# Lysine acetylation plays a role in RNA binding protein-regulated alternative pre-mRNA splicing

**DOI:** 10.1101/2025.07.11.664476

**Authors:** Christopher Nunez, Gustavo Salgado, UyenPhuong Tran, Azeem Horani, Samantha Dreyer, Tyler Luchko, Niroshika M. Keppetipola

## Abstract

Alternative pre-mRNA splicing allows one gene to encode multiple spliced messenger RNAs and, in turn, multiple proteins from a single gene transcript. This process is tightly regulated by cis elements within the pre-mRNA and trans-acting RNA binding proteins that recognize and bind to these elements, thus influencing the spliceosome assembly at adjacent splice sites. Thus, chemical modifications in either the cis-elements or trans factors or both can significantly alter splicing patterns and, thereby, the cellular proteome. Recent studies highlight that many RNA binding proteins (RBPs) are modified at multiple lysine side chains via acetylation, which neutralizes the formal positive charge and disrupts RPBs’ ability to participate in RNA recognition, binding and protein-protein interactions. This suggests that lysine acetylation of RPBs may be a novel mode of eukaryotic gene regulation during pre-mRNA processing. To test this, we used the well-characterized polypyrimidine tract binding protein (which is acetylated at several lysine side chains) as a model system to investigate the role of reversible RBP acetylation in regulating alternative-pre mRNA splicing. Using multiple sequence analysis, structure-based electrostatic modeling of RNA-protein interactions, and multi-site glutamine (acetyllysine mimic) and arginine (deacetyllysine mimic) mutants, we show for the first time that for a subset of PTBP1-regulated exons, acetylation at RNA-interacting lysine side chains significantly alters PTBP1 splicing activity.

## Introduction

Alternative splicing of pre-mRNA transcripts is an important step in eukaryotic gene expression that contributes to a diverse transcriptome and in turn, a diverse proteome (Black 2003; Matlin, Clark, and Smith 2005; Chen and Manley 2009; Lee and Rio 2015). This process is regulated in part by trans-acting RNA binding proteins that bind to corresponding cis-elements and influence the assembly of a functional spliceosome at adjacent splice sites to either enhance or repress the inclusion of a regulated exon. Thus, factors that influence the amounts and activity of RNA binding proteins dictate the cellular transcriptome and in turn the proteome.

Recent studies highlight that many RNA binding proteins (RBPs) are acetylated at lysine side chains.(Narita, Weinert, and Choudhary 2019; Choudhary et al. 2009a; Kim et al. 2006; Zhang et al. 123AD; Cohen et al. 2015a; Babic, Jakymiw, and Fujita 2004a; Sasaki et al. 2012a; Arenas et al. 2020a; Gal et al. 2019a; Pina et al. 2018a; Hornbeck et al. 2015a; Castello et al. 2016; Sullivan et al. 2025). The positively charged **ε**-amino group of lysine side chains can participate in electrostatic interactions including salt bridges, hydrogen bonds and stacking interactions with nitrogenous bases in the RNA molecule. Such interactions are frequently found at RNA-protein interfaces.(Corley, Burns, and Yeo 2020; Snead and Gladfelter 2019) Acetylation at the **ε**-amino group neutralizes the formal positive charge and renders a delocalized lone pair on the nitrogen atom (by participating in resonance with the adjacent carbonyl of the acetyl group). Thus, acetylation significantly decreases the degree of positive charge on lysine side chains, which can reduce RBP capacity to interact with the negatively charged phosphodiester backbone, its interaction strength with the aromatic rings of the nitrogenous bases via pi-cation interactions, and its ability to act as an H-bond acceptor. (Fig. 1).

**Figure 1.**
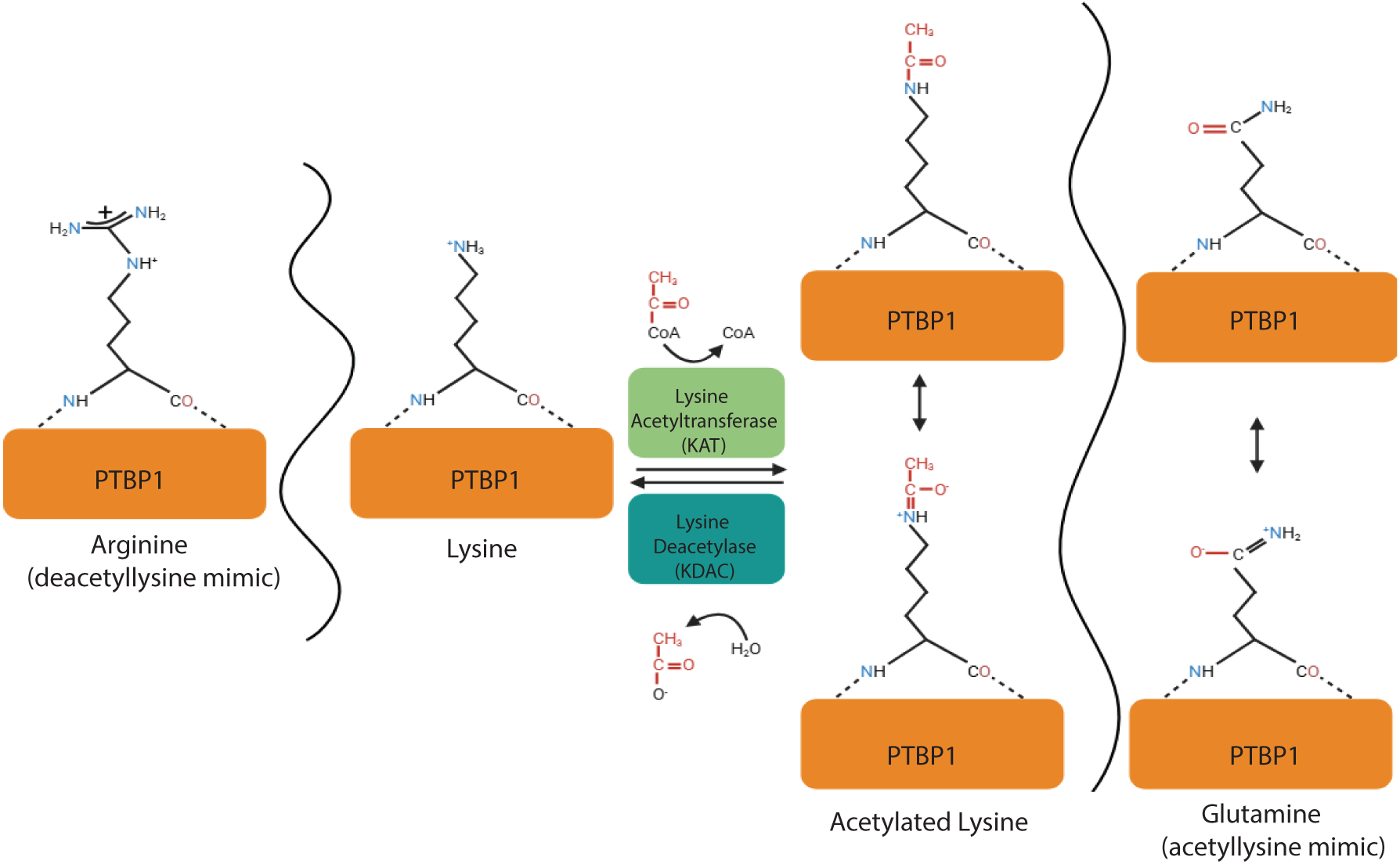
Side chain charge distribution of lysine, glutamine (acetylated lysine mimic) and arginine (deacetylated lysine mimic). **Center:** Lysine acetyltransferase (KAT) catalyzes the addition of an acetyl group using acetyl-CoA as a cofactor. A lysine deacetylase (KDAC) catalyzes the removal and release of the acetyl group. Resonance structures of the acetylated lysine side chain is indicated. Acetyllysine-mimic glutamine side chains’ electron delocalization is indicated on the right. Deacetyllysine mimic arginine side chain electron delocalization is indicated on the left.

It is plausible that reversible lysine acetylation can modulate the activity of RNA binding proteins (including recognition and binding to cis-elements and interacting with spliceosome components) to dictate distinct alternative splicing patterns and, in turn, a corresponding cellular proteome. For example, lysine acetylation-deacetylation plays a critical role in regulating transcription machinery in the context of histone tails, including RNA polymerase access to the genes and transcription (Ghoneim, Fuchs, and Musselman 2021; Anderson, Lowary, and Widom 2001). Similarly, a lysine acetylome study revealed RNA binding domains are the most significantly overrepresented domain architecture of acetylated proteins (Choudhary et al. 2009b). Moreover, the study also showed proteins involved in RNA splicing as the second most abundant class of acetylated proteins next to those involved in the cell cycle supporting a yet-to-discovered role for RBP acetylation. Furthermore, the study revealed more acetylation sites than acetylated proteins involved in splicing, indicating that multiple lysine residues are acetylated on a single protein. RNA binding was also reported as the most significantly enriched molecular function in another lysine acetylome study with 85 out of 184 acetylated proteins playing a role in the spliceosome and RNA transport.(Zhang et al. 123AD). Prompted by these findings, here, we aimed to test the role of lysine acetylation in RBP-regulated alternative pre-mRNA splicing: a novel mode of eukaryotic gene regulation at the level of pre-mRNA processing.

To this end, we utilized polypyrimidine tract binding protein 1 (PTBP1) as a model system.(Romanelli, Diani, and Lievens 2013; Niroshika Keppetipola et al. 2012) PTBP1 is a well-characterized RNA-binding protein involved in alternative splicing regulation (Chan and Black 1997; Sharma, Falick, and Black 2005a; 2005b; N Keppetipola et al. 2012; Sharma et al. 2014a; 2008a; Wongpalee et al. 2016; N. M. Keppetipola et al. 2016; Ontiveros et al. 2020). PTBP1 is composed of four RNA binding domains (RBDs), connected by three linker regions and an N-terminal region. PTBP1 most often functions to repress inclusion of regulated exons in spliced mRNA but can enhance their inclusion as well.(Boutz et al. 2007; Wollerton et al. 2001; Llorian et al. 2010; Spellman et al. 2005; Xue et al. 2009). Solution structures of each PTBP RBD bound to an RNA hexamer reveal residues that play a critical role in PTBP1 RNA binding (Oberstrass et al. 2005). We discovered that PTBP1 is acetylated in both non-neuronal HeLa and neuronal WERI nuclear extracts.(Pina et al. 2018b; Sullivan et al. 2025) Our results indicate that PTBP1 is acetylated at lysine side chains (many of which make direct contact with the RNA) in all four RNA-binding domains. Moreover, discovery-based proteomics conducted by others also revealed acetyl modifications on some of these RNA-binding lysine residues (Hornbeck et al. 2015b). The mechanism of PTBP1-mediated splicing regulation is well understood.(Sharma, Falick, and Black 2005c; Sharma et al. 2014b; 2011; 2008b; Clerte and Hall 2009a; Auweter, Oberstrass, and Allain 2007; Lamichhane et al. 2010a) Thus, we deemed PTBP1 an ideal candidate to test the role of RBP acetylation in alternative splicing regulation.

PTBP1 RBDs 3 and 4 comprise the minimal region required to bind to single-stranded RNA found in splicing regulatory cis-elements (Clerte and Hall 2009b). Moreover, the RBD 3 and 4 regions have a unique domain arrangement that can induce RNA looping, which is suggested to play a role in PTBP1-mediated exon repression.(Lamichhane et al. 2010b; Kafasla et al. 2012; Vitali et al. 2006; Oberstrass et al. 2005). PTBP1 RBD3 and 4 have five and three acetylated residues, respectively (Pina et al. 2018b; Hornbeck et al. 2015b; Sullivan et al. 2025).

In this study, we sought to address the role of lysine acetylation in RBD 3 and 4 in PTBP1-regulated splicing repression activity. To this end, we assayed single alanine mutants, multi-site glutamine (acetyllysine mimic), and multi-site arginine (deacetyllysine mimic) mutants for splicing activity with three PTBP1-regulated exons. We modeled changes in the ion distribution using the 3D reference interaction site model (3D-RISM)(Kovalenko and Hirata 1999) of molecular solvation to investigate electrostatic changes in the RNA interaction surface of RBDs 3 and 4 upon acetylation. Our results reveal a novel role for acetylation in PTBP1-regulated splicing repression. Importantly, our findings validate RBP acetylation as a novel mode of gene regulation in eukaryotes.

### PTBP1 acetylated lysine residues are evolutionarily conserved

We conducted a multiple sequence alignment to assess the conservation of the lysine residues acetylated in the RBD3 and RBD4 regions of the PTBP1 protein across different species (Fig. 2A and 2B). We used PTBP1 isoform 4 (PTBP1.4) in our study and residue numbers correspond to PTBP1.4 (hereafter referred to as PTBP1). Our data highlight that Lys394, Lys400, and Lys436 in RBD3, and Lys508 and Lys511 in RBD4 are invariant, indicating an important role in PTBP1 function. Lys424, Lys454 and Lys515 are less conserved, with some species having a conservative arginine substitution, highlighting an important role for a positive charge at these positions. Lys424 participates in a salt bridge interaction with Glu528, which plays a critical role in maintaining the unique fold between RBDs3 and 4 (Fig. 3). Our data highlight that a positive charge at position 424 is sufficient to maintain this interaction. Collectively, our results highlight an important role for these lysine residues in PTBP1 function.

**Figure 2.**
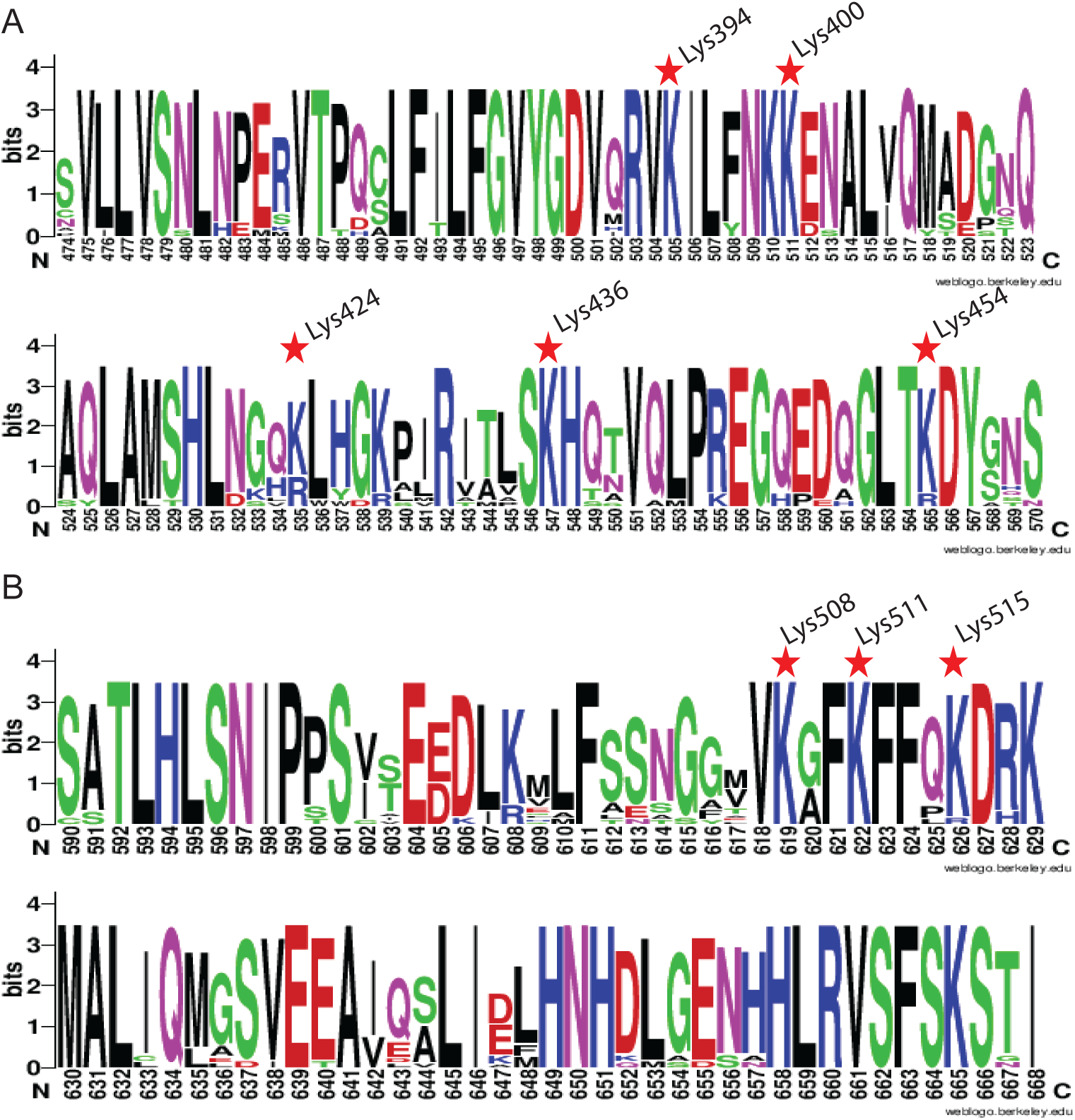
Weblogo of PTBP1 (A) RRM3 and (B) RRM4 showing the high degree of sequence conservation. MSA were created using ClustalW (as implemented in MEGA v11) from 17 type of species (Humans, Chimpanzee, Mouse, Alpine Marmot, Little Egret, Red Junglefowl, Ruddy Duck, African Clawed Frog, Rhinatrema, Tiger Snake, Snapping Turtle, Bynoes Gecko, Zebrafish, Chinese Tongue Solefish, Lung Fish, Fruit Fly, and Roundworm). The highly conserved lysine residues known to be acetylated in splicing reaction conditions are highlighted with a red star and residue number.

**Figure 3.**
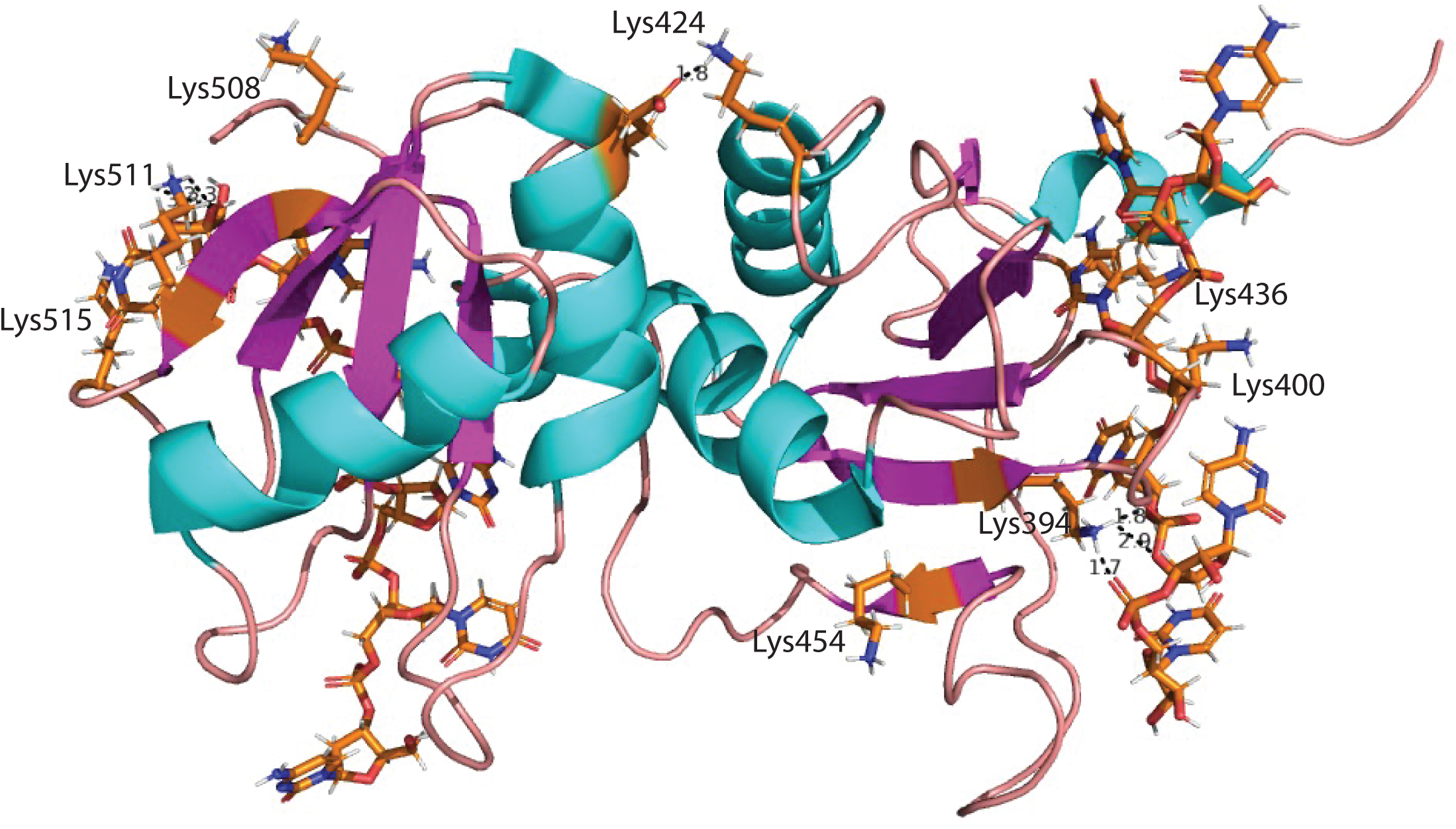
Location and interactions of modified lysine side chains in RRMs 3 and 4. Cartoon representation of NMR solution structure of PTBP1 RRM3_4 (2ADB) (Oberstrass et al. 2005). The main chain cartoon traces are colored by secondary structure with alpha helices in cyan, beta sheet in magenta and turns and loops in pink. Post-translationally acetylated residues identified by mass spectrometry are labeled. Side chains of modified residues and the CUCUCU RNA hexamer (for PTBP1) are shown as sticks and colored by element.

### Acetylated lysine side chains interact with RNA

Here, we used the available solution NMR structure of the PTBP1 RBD3 and 4 domains bound to a CUCUCU hexamer to determine the position of acetylated lysine residues and specific interactions between the side chains and the RNA (Fig. 3) (Oberstrass et al. 2005). We note Lys394 makes two H-bond interactions with the phosphodiester backbone between C5 and U6 and makes another H-bond with the terminal 5’phosphate of U6. Thus, Lys394 plays an important role in binding affinity and orienting the bound substrate. Lys400 does not make H-bond interactions with the substrate in this structural snapshot yet has the potential to do so given its location in a flexible loop region. Lys424 participates in a salt-bridge interaction (with Glu528) that is critical for maintaining the unique arrangement between RBDs 3 and 4. It is plausible that a positively charged arginine side chain can substitute for the lysine side chain as highlighted in the sequence conservation (Fig. 2). Lys436 side chain is optimally positioned, underneath the nitrogenous base of C3, and can participate in pi-cation interactions stabilizing RNA binding. Lys454 is positioned in a flexible loop region that can likely participate in RNA binding with substrates longer than six nucleotides. Thus, acetyl modifications at these side chains can disrupt RBD3-RNA non-covalent interactions that contribute to binding affinity.

In the RBD4 region, Lys508 has the potential to participate in RNA binding with longer RNA sequences. The lysine side chain of Lys511 contributes two H-bond donors to form H-bonds with 3’OH (H-bond acceptor) of U6. Lys515 is ideally positioned to interact with the nitrogenous base of U6 via a pi-cation interaction. We surmise either lysine or arginine can participate in this interaction, which is consistent with the conservative arginine substitution at this position (Fig. 2). Thus, acetylation at lysine side chains in RBDs 3 and 4 can disrupt salt bridges, H-bond interactions, decrease the strength of pi-cation interactions, and introduce steric bulk that collectively can modulate PTBP1 RNA binding affinity and in turn splicing activity.

### Acetylation alters local ion concentrations and disrupts RNA binding

We carried out 3D-RISM calculations to investigate how RBD3/4 acetylation might influence local ion concentration. Acetylation of lysine residues leads to significant changes in the RNA-binding affinity of the RBD3/4 domains. These changes are due to both direct alterations in protein–RNA interactions and broader effects on the surrounding ion distributions.

As illustrated in Figure 4, in the wild-type holo state (PDB ID: 2ADC), three lysine residues in RBD3 form salt bridges with the RNA phosphate backbone, helping to maintain the RNA in a bound state. Similar structure and interactions are also observed in RBD4 (not shown). Upon acetylation, mimicked by lysine-to-glutamine mutations, these salt bridges are lost and replaced with weaker hydrogen bonds between the RNA and protein side chains. Additionally, the loss of charge at these sites may lead to conformational changes in RBD3/4, which may disrupt the structural integrity of the RNA-binding channel. In the apo state (PDB ID: 2EVZ), where RNA is not bound, these lysine residues show greater separation, further supporting their role in RNA stabilization.

**Figure 4:**
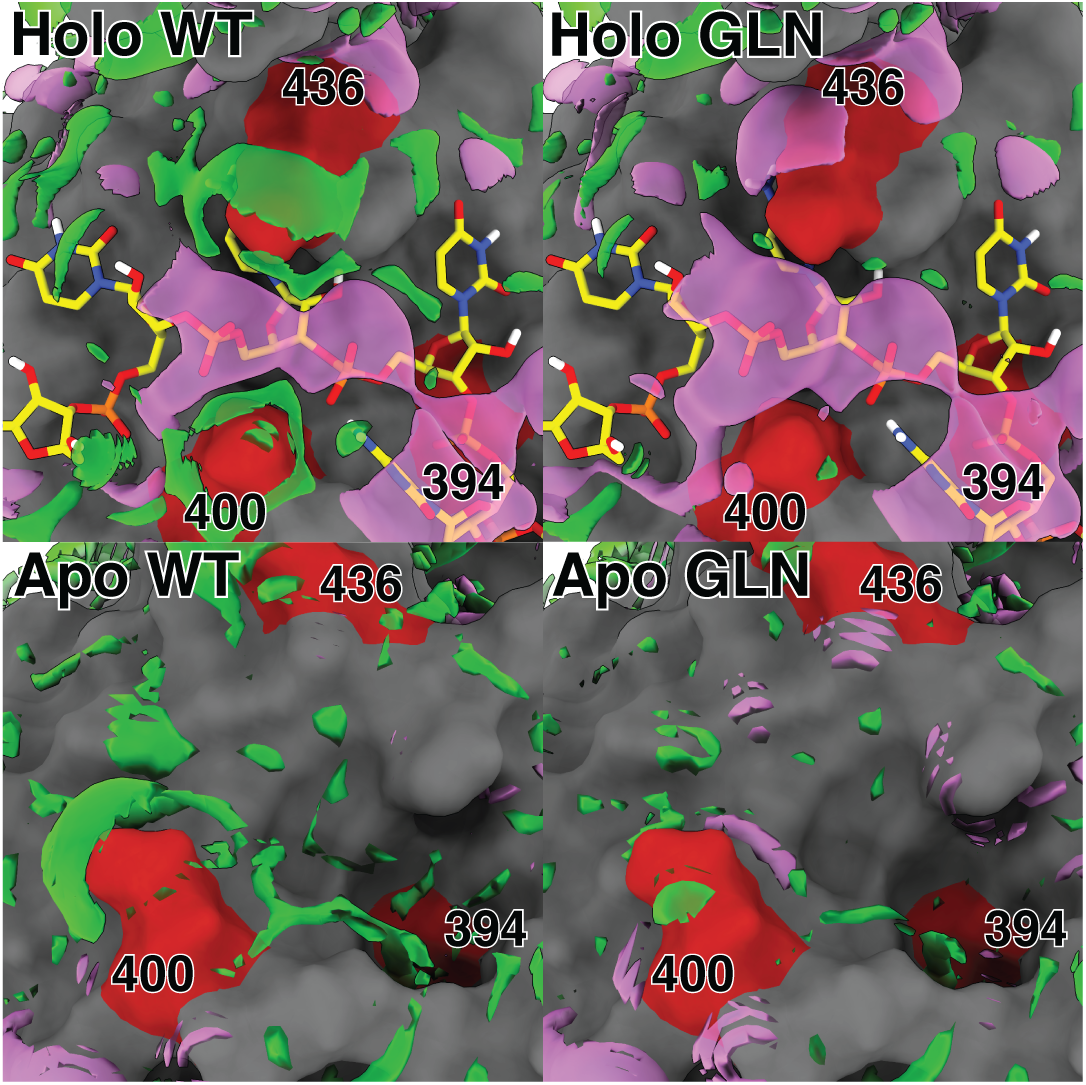
Ion distributions around the RNA-binding site of holo (top, PDB ID: 2ADC) and apo (bottom, PDB ID: 2EVZ) RBD3. Right: unacetylated lysine is positively charged, forming salt bridges with the RNA backbone and drawing in excess chloride ions. Left: glutamine is substituted for lysine, mimicking acetylation and neutralizing the charge. This weakens binding by removing the salt bridges and shifts the local ion concentration to contain more potassium and less chloride. Potassium (violet) and chloride (green) distributions are shown at 0.8 M isosurfaces (four times bulk concentration). RNA is shown as sticks (carbon: yellow, nitrogen: blue, oxygen: red, hydrogen: white, phosphorus: orange), RBD3 as grey solvent accessible surface, and lysine residue 394, 400 and 436 as red solvent accessible surfaces.

The mutations that mimic acetylation also significantly affect ion distributions around RBD3/4. In the holo state, the net charge changes from -3e in the wild-type to -11e in the mutant, while for the apo state, the net charge shifts from +7e to -1e. These changes in net charge are reflected in the excess ion distributions, as excess potassium increases and chloride decreases (Table 1). The increase in potassium ions around the protein suggests competition with the RNA phosphate groups for electrostatic interactions, which could further reduce protein–RNA affinity.

**Table 1.** Excess number of potassium and chloride ions around wild-type and mutated apo and holo structures of PTBP1 RDB 3 and 4.

| Structure | Excess K<sup>+</sup> | Excess Cl<sup>-</sup> |
| --- | --- | --- |
| WT Holo (PDBID: 2ADC) | -3.1 | -6.1 |
| Mutant Holo (PDBID: 2ADC) | +1.4 | -9.6 |
| WT Apo (PDBID: 2EVZ) | -7.4 | -0.4 |
| Mutant Apo (PDBID: 2EVZ) | -3.5 | -4.5 |

These shifts are evident within the RNA-binding channel itself. In the wild-type holo state, chloride ions cluster around the lysine residues, and potassium follows the RNA backbone (Fig. 4) (Giambaşu et al. 2014). In contrast, the glutamine mutant (acetylation mimic) shows a loss of chloride accumulation and an expanded high-density potassium region around the RNA. Notably, the wild-type apo state has no regions where the potassium concentration exceeds 0.8 M in the binding channel, whereas small potassium pockets appear in the mutant, likely competing with the protein for RNA binding. Together, these findings demonstrate that acetylation, by removing salt bridges and altering local electrostatic environments, weakens RNA binding. The effect arises both from local disruption of lysine–RNA interactions and from global changes in net charge and ion distributions that collectively reduce electrostatic attraction between the RNA and RBDs 3 and 4.

### Single lysine-to-alanine mutants do not affect PTBP1 splicing activity on the regulated Dup175 test exon

We created single lysine-to-alanine mutations of residues acetylated in RBDs 3 and 4 and assayed these mutants for protein expression and splicing activity on the PTBP1-regulated Dup175 test exon (Fig. 5). The Dup175-DS9 minigene (Fig. 5A) encodes a three-exon pre-mRNA, where the central test exon is regulated by two high-affinity PTBP1 binding sites, one upstream of the branch point sequence and the other within the test exon itself. Previous studies highlight that this exon is repressed by PTBP1 (Amir-Ahmady et al. 2005).

**Figure 5.**
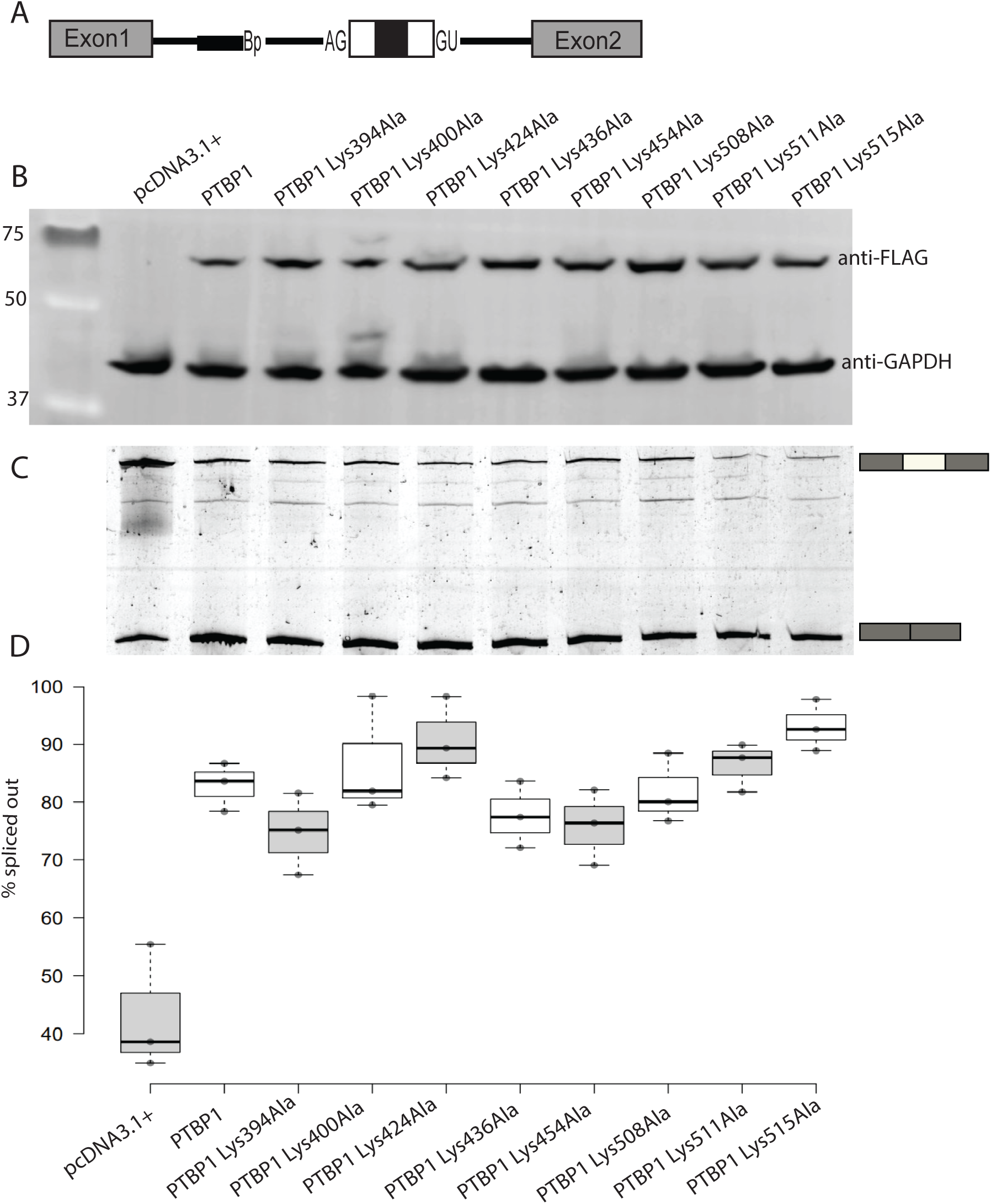
Single-point mutations of acetylated lysine residues do not play a role in PTBP1 splicing activity. **(A)** Structure of reporter minigene Dup175-DS9, with β-globin exons 1 and 2 flanking the hybrid exon and its intronic regulatory regions. The test exon is flanked by PTBP binding sites from Src and by wild-type β globin exons 1 and 2 (Dominski and Kole 1991b; Modafferi and Black 1997; Amir-Ahmady et al. 2005). (B) Immunoblot of Flag-PTBP1.4 protein and RBD3- and RBD4-alanine mutants in cell lysates after transfection with 2.0 µg of Flag-PTBP plasmid DNA. Protein expression levels were quantified and normalized to GAPDH. (C) Splicing reporter Dup 175-DS9 was cotransfected with empty expression vector (pcDNA 3.1 +) or PTBP1 wild-type or mutant expression plasmid (2.0 µg). RNA was harvested after 48 hours, assayed by RT-PCR, and quantified. Lanes are aligned with labeled lanes in panel B. (D) To quantify the level of repression, percent-spliced-out (PSO) was calculated by dividing band intensity for excluded product by the total value of excluded and included products. The PSI values (n=3) were used to generate the boxplots using BoxPlotR code (Spitzer et al. 2014). Center lines show the medians; box limits indicate the 25^th^ and 75^th^ percentiles as determined by R software; data points are plotted as circles. Statistical significance was determined by one-way Analysis of Variance (ANOVA), followed by a post hoc Tukey Honestly Significant Difference (HSD) test.

Our data reveal that all single alanine mutants are well expressed (Fig. 5B). Wild-type PTBP1 repressed the Dup175 exon by 82.9% (Fig. 5C and D). Our splicing data highlight that the absence of a positively charged lysine side chain at one of the eight positions (394, 400, 424, 436, 454, 508, and 511) did not alter PTBP1 splicing repression activity significantly. Thus, data from this experiment highlight that removing a single lysine side chain (and in turn a positive charge) does not exert significant changes in PTBP1 splicing activity. Given the multiple contacts made by PTBP1 RBD3 and 4 with RNA, we surmised a multi-site coordinated model for acetylation-regulated PTBP1 splicing activity.

### Multi-site Gln mutations in PTBP1 RBD3 and RBD4 play a role in PTBP1-regulated splicing repression of the neuronal cSrc N1 exon

We used multi-site domain-specific mutants that mimic acetylated versus deacetylated states of RBD3, RBD4, and combined RBD3-RBD4. To this end, we used multi-site lysine-to-glutamine mutants to mimic the acetylated lysine side chains in RBD3 (K394Q-K400Q-K424Q-K436Q-K454Q) and the RBD4 region (K508Q-K511Q-K515Q). We also generated counterpart multi-site arginine mutants that mimic deacetylated lysine side chains (K394R-K400R-K424R-K436R-K454R and K508R-K511R-K515R). These residues are not modified by acetylation and retain the formal positive charge. Given the unique domain organization, we also created multi-site glutamine and arginine mutants combined in the RBD3-RBD4 region to determine whether acetylation-deacetylation might exert splicing effects cooperatively. We note these multi-site mutants mimic two distinct states of the PTBP1 protein, where either the RBD3, RBD4 or RBD3-4 is fully acetylated or deacetylated under experimental conditions. This is in contrast to wild-type PTBP1, which is subject to dynamic changes in the extent and degree of acetylation.

We assayed these mutants for protein expression and splicing activity in vivo on the PTBP1-reglated N1 exon utilizing the Dup4-5 splicing reporter minigene (Fig. 6A) (Modafferi and Black 1999). PTBP1 represses the inclusion of the N1 exon in spliced mRNA. The central exon of this minigene encodes the endogenous neural-specific N1 exon and is flanked by portions of the endogenous introns. Western blot data reveal that these mutants are well expressed (Fig. 6B). The splicing data highlight that PTBP1 represses inclusion of the N1 exon as previously reported. We obtained a percent spliced in (PSI) value of 2.5 for wild type PTBP1. Our data reveal that multi-Gln RBD3 mutant, which mimics an acetylated RBD3, increased PSI to 9.1 (thereby decreasing repression) and this is significantly different to wild type PTBP1 (p =0.0001). Thus, our data demonstrate a role for the five acetylated lysine residues in RBD3 in PTBP1 splicing activity. The counterpart multi-Arg RBD3 mutant demonstrated a PSI of 2.5, similar to wild-type PTBP1, signifying a role for electrostatic interactions exerted by positively charged side chains in splicing repression activity. The RBD3-4 multi-Gln yielded a PSI of 16.9, significantly higher than wild type PTBP1 (p<0.00005). Thus, our data highlight that lysine acetylation of eight residues in RBD 3 and 4 further decreases PTBP1 splicing repression activity. The counterpart RBD3-4 Arg mutant demonstrated a PSI of 3.9, which is not significantly different from wild-type PTBP1. Thus, our data support that positively charged side chains at these positions restore splicing repression to wild-type PTBP1 levels. The RBD4 multi-Gln mutant and the counterpart RBD4 multi-Arg mutant showed a PSI of 3.8 and 3.7, respectively. These values are not significantly different from wild-type PTBP1.

**Figure 6.**
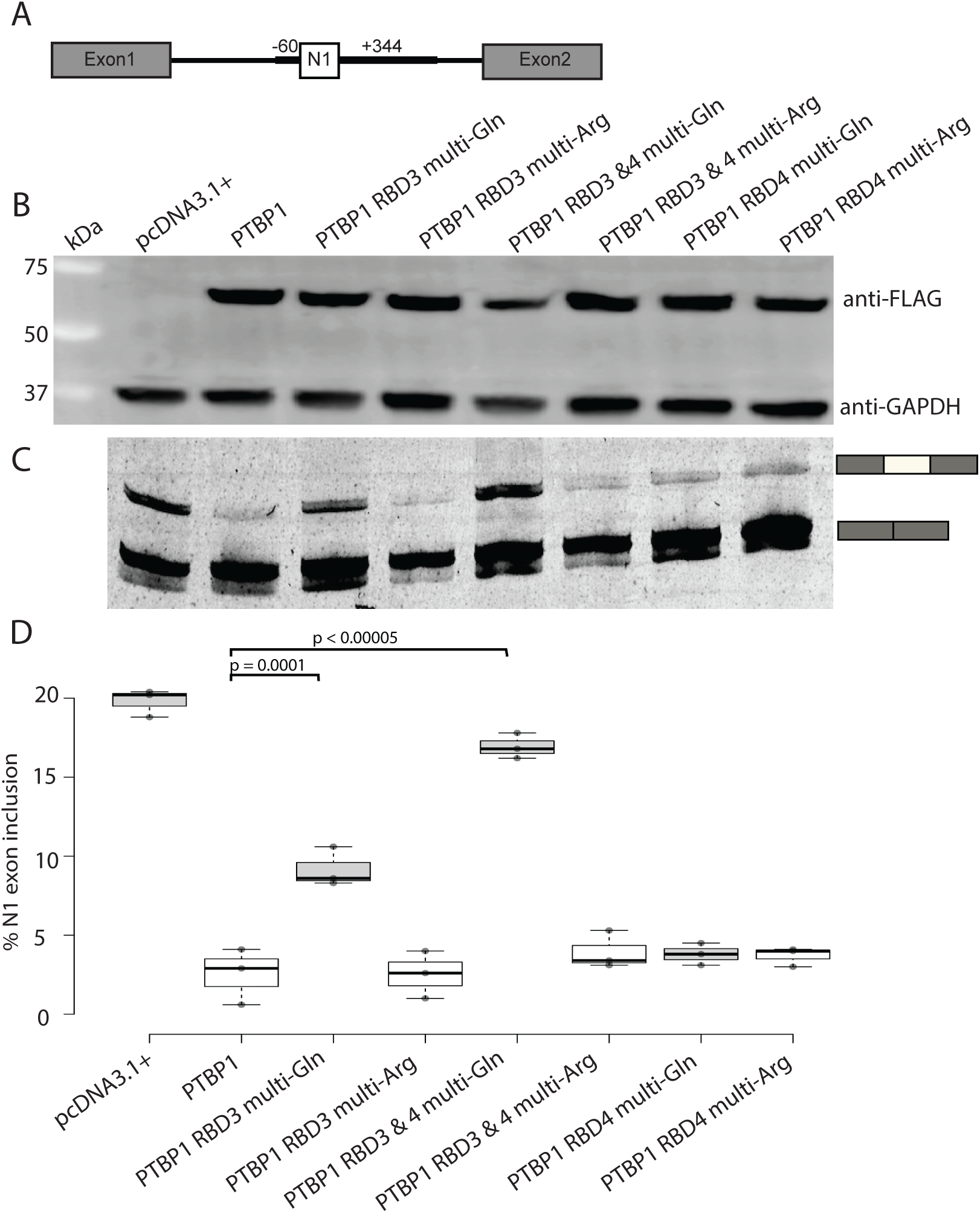
Acetylation in PTBP1 RBD3 and RBD4 plays a role in N1 exon alternative splicing. (A) Structure of reporter minigene Dup 4-5 N1 exon, with β-globin exons 1 and 2 flanking the exon and its intronic regulatory regions. (B) Immunoblot of Flag-PTBP1.4 protein and RBD3, RBD4, and RBD3-4 acetyllysine- and deacetyllysine-mimic mutants in cell lysates after transfection with 2.0 µg of Flag-PTBP plasmid DNA. Expression levels were visualized using anti-Flag antibody, and GAPDH served as a loading control. (C) Splicing of the c-Src N1 exon by PTBP1.4, acetyl, and deacetyl mimic mutants. Dup4-5 (0.5µg) was co-transfected with empty vector pcDNA 3.1 (+) or PTBP1 wild-type and mutant expression plasmids (2.0 µg). RNA was harvested after 48 hours, assayed by RT-PCR, and quantified. Lanes are aligned with labeled lanes in panel B. (D) The level of PSI was calculated by dividing band intensity for the included product by the total value of excluded and included products. The PSI values (n=3) were used to generate the boxplots using BoxPlotR code (Spitzer et al. 2014). Center lines show the medians; box limits indicate the 25^th^ and 75^th^ percentiles as determined by R software; data points are plotted as circles. Significant p-values are presented above the plot.

These findings suggest that lysine acetylation in RBD4 (in the absence of acetylation in RBD3) does not play a role in PTBP1-regulated N1 exon splicing repression. We note the decrease in splicing repression activity is approximately two-fold greater for the RBD3-4 multi-Gln (PSI 16.9) than in the RBD 3 multi-Gln mutant (PSI 9.1). Thus, our results suggest lysine acetylation in RBD3 plays a primary role in mediating the observed changes and that acetylation in RBD4 works cooperatively to enhance this effect. This result agrees with previous studies that show RBD3 is the major determinant of binding affinity and specificity, and further supports the role that residues outside RBD3, including RBD4, play in electrostatic interactions in binding and complex stabilization (Perez and Patton 1997).

### Multi-site Gln mutations in PTBP1 RBD3 and RBD4 play a role in PTBP1-regulated splicing repression of the calcium channel Exon 8a

We next assayed these multi-site Gln and Arg mutants in RBD3, RBD4 and RBD3-4 for splicing activity in vivo on the PTBP1-regulated exon 8a from the *CACNA1C* gene. PTBP1 represses inclusion of exon 8a (E8a) of the CaV1.2 calcium channel transcript. The Dup4-1 E8a splicing reporter minigene contains the same basic components as Dup175-DS9. However, it encodes exon 8a (in lieu of the Dup175 central test exon) and flanking intron sequences from the original mouse gene (Fig. 7A). The intron sequence upstream of E8a contains several CU-rich elements known to be high-affinity PTBP1 binding sites (Tang et al. 2011). Western blot data indicate that the mutants are well expressed (Fig. 7B). Splicing data demonstrate that PTBP1 represses inclusion of exon 8a with a PSI of 27.3 (Fig. 7C and D). Our data reveal that the multi-Gln RBD3 mutant, which mimics an acetylated RBD3, decreased repression (increasing inclusion) to a PSI of 65.8 and is significantly different from wild type PTBP1 (p =0.001). Multi-Arg RBD3 restored splicing activity to wild-type PTBP1 levels (PSI of 30.7). Thus, our data support a role for positive charges at these five positions in RBD3 for PTBP1 splicing repression of exon 8a. Our data also reveal glutamine side chains at these five positions, which mimic acetylation, can significantly alter PTBP1 splicing activity. A multi-Gln RBD3-4 mutant further decreased splicing repression to a PSI of 78.4 and is significantly different from wild-type activity (p<0.00005). Multi-Arg RBD3-4 mutants restored splicing repression to wild-type levels with a PSI of 30.5.

**Figure 7.**
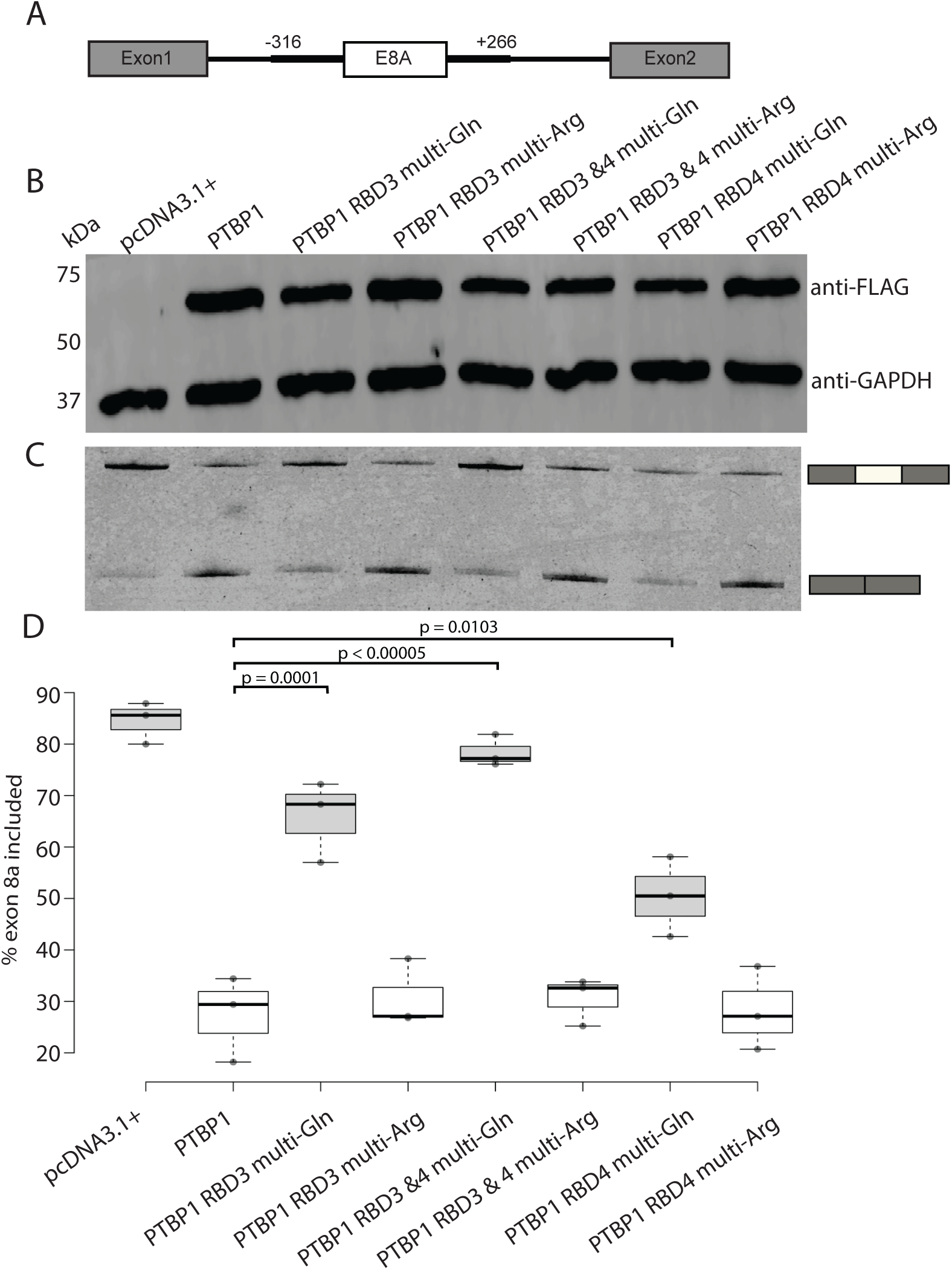
Acetylation in PTBP1 RBD3 and RBD4 plays a role in calcium channel exon 8a alternative splicing. (A) Structure of reporter minigene Dup 4-1 E8a, with β-globin exons 1 and 2 flanking the regulated exon 8a and its intronic regulatory regions (Dominski and Kole 1991a; Modafferi and Black 1997). (B) Immunoblot of Flag-PTBP1.4 protein and RBD3, RBD4, and RBD3-4 acetyllysine- and deacetyllysine-mimic mutants in cell lysates after transfection with 2.0 µg of Flag-PTBP plasmid DNA. Expression levels were visualized using anti-Flag antibody, and GAPDH served as a loading control. (C) Splicing of exon 8a by PTBP1.4, acetyl, and deacetyl mimic mutants. Dup4-1 E8a (0.5µg) was co-transfected with empty vector pcDNA 3.1 (+) or PTBP1 wild-type and mutant expression plasmids (2.0 µg). RNA was harvested after 48 hours, assayed by RT-PCR, and quantified. Lanes are aligned with labeled lanes in panel B. (D) The level of PSI was calculated by dividing band intensity for the included product by the total value of excluded and included products. The PSI values (n=3) were used to generate the boxplots using BoxPlotR code (Spitzer et al. 2014). Center lines show the medians; box limits indicate the 25^th^ and 75^th^ percentiles as determined by R software; data points are plotted as circles. Statistical significance was determined by one-way ANOVA, followed by a post hoc Tukey HSD test. Significant p-values are presented above the plot.

Thus, our findings reveal that positive charges at the five RBD3 and three RBD4 positions play a significant role in PTBP1-mediated exon 8a splicing repression. Our data indicate that acetylation at these side chains significantly reduce repression and thereby modulate PTBP1-regulated splicing of exon 8a. Multi-Gln RBD4 decreased splicing activity to a PSI of 50.4, which is significantly different from wild-type PTBP1 (p=0.0103). We note this value is lower than PSIs for multi-Gln RBD3 (PSI 65.8) and multi-Gln RBD3-4 mutants (PSI 78.4). Multi-Arg RBD4 demonstrated a PSI of 28.2, similar to wild-type PTBP1 splicing activity.

Collectively, data from this experiment further support a role for the positively-charged lysine side chain at five and three positions in RBD3 and RBD4, respectively, in PTBP1 splicing activity. Notably, our findings highlight that removing the formal positive charge via acetylation alters electrostatic interactions, thereby modulating PTBP1-regulated exon 8a splicing activity.

## Discussion

To date, only a few studies have highlighted the role of lysine acetylation in RBP function by altering RBP-protein, RBP-RNA interactions, and subcellular localization.(Cohen et al. 2015b; Babic, Jakymiw, and Fujita 2004b; Sasaki et al. 2012b; Arenas et al. 2020b; Gal et al. 2019b). In this study, we report for the first time a role for lysine acetylation in RBP-regulated pre-mRNA splicing; acetylation significantly decreases PTBP1-mediated splicing repression of the neuronal N1 exon and the calcium channel exon 8a (Fig. 8). Ongoing experiments in our laboratory aim to determine whether this decrease in repression is mediated via changes in RNA binding affinity and/or protein-protein interactions. Our data indicate that acetylation of multiple lysine residues within RBD3 and RBD4 are necessary for regulation of the N1 exon and calcium channel exon 8a. This result is consistent with reports indicating multiple acetylation sites in RNA binding proteins (Choudhary et al. 2009b).

**Figure 8.**
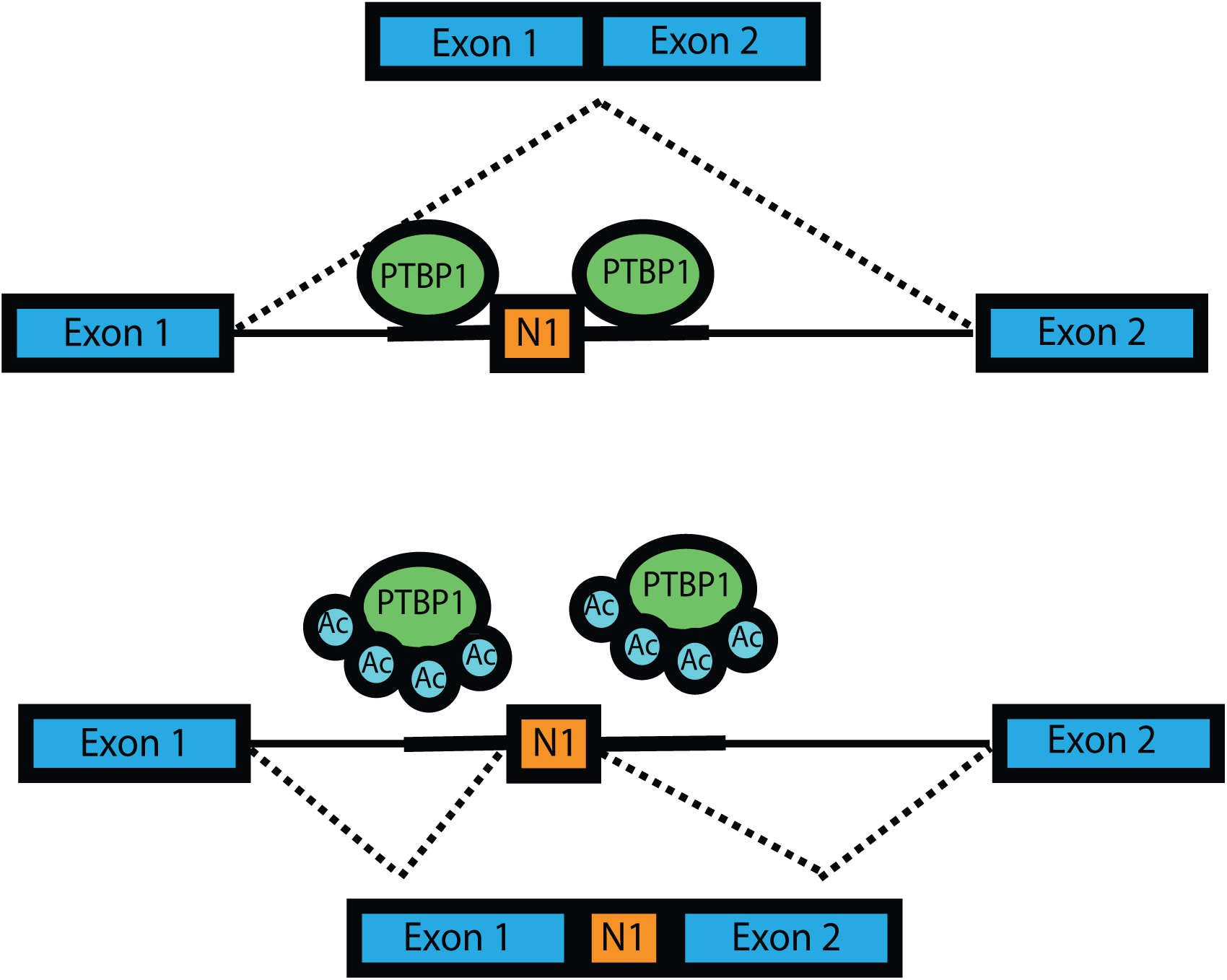
Model for lysine acetylation modulation of PTBP1-regulated splicing of the N1 exon. Lysine acetylation significantly decreases PTBP1-mediated splicing repression of the neuronal N1 exon.

Lysine acetyltransferases (KATs) and lysine deacetylases catalyze the addition and removal of acetyl groups from lysine side chains. KATs primarily use the cellular metabolite acetyl-CoA as a cofactor when catalyzing acetylation.(Sheikh and Akhtar 2019; Shvedunova and Akhtar 2022). Thus, the cellular concentration of acetyl-CoA plays a major role in determining the targets and kinetics of acetylation.(Shvedunova and Akhtar 2022) Therefore, RBPs, including PTBP1, that undergo acetylation can serve as key effectors in integrating the cellular metabolic state with alternative splicing patterns and gene expression. Moreover, PTBP1 acetylated lysine side chains are also methylated, highlighting possible crosstalk between acetylation and methylation in response to cell-signaling and metabolic state (Sullivan et al. 2025). Our previous work has shown that PTBP1 has post-translational modifications, including phosphorylation, acetylation, and methylation (Pina et al. 2018b; Sullivan et al. 2025). Yet, the role of acetyl and methyl modifications in RBP function is not well understood. Similar to DNA-binding histone proteins, we propose a scenario for post-translational modifications of RNA-binding proteins as a reversible “on-off switch” brought about by a set of enzymes that are part of signaling pathways in response to stimuli, including the cellular metabolic state. Ongoing and future work aims to identify writers, erasers, and readers of these modifications and their role in mediating RBP-regulated alternative splicing.

In sum, our findings show a novel role for acetylation in PTBP1-regulated alternative pre-mRNA splicing with certain exons. Moreover, our results reveal a previously unrecognized role for post-translational acetylation in PTBP1-mediated splicing regulation, introducing a new paradigm in eukaryotic gene regulation. Given the prevalence of acetylation in RNA-binding proteins and their role in regulating alternative pre-mRNA splicing, our results indicate a novel mechanism of gene regulation in eukaryotes that warrants further investigation.

## Materials and Methods

### Cell culture and transfections

Mouse neuro 2A (N2A) cells were grown according to the American Type Culture Collection (ATCC) recommended protocols in DMEM (Fisher Scientific) with 10% FBS (Omega Scientific) and 1X PenStrep Glutamine (Life Technologies). Each experiment was conducted thawing freezes to ensure low passage cells and passage #3 was routinely used for assays to maintain consistency across experiments. Cell morphology was frequently checked during subculturing to identify unusual appearance and if observed, a new batch of frozen cells were cultured. Transfections were carried out using Lipofectamine 3000 transfection reagent (Life Technologies) according to the manufacturer’s instructions. Cells were harvested 48 hours post-transfection and RNA and protein were isolated for analysis.

### Plasmid Construction

Single alanine mutants were created by two-step overlap extension PCR. Primers were designed to introduce single mutations at corresponding sites, carrying overlapping flanking sequences. The PCR fragments were cloned into pcDNA3.1(+) (Life Technologies) using its BamH1 and EcoRV restriction sites. The inserts were sequenced to verify the absence of unwanted coding changes during PCR amplification. Each expression plasmid carried an N-terminal FLAG-tag. The minigene reporter Dup175-DS9 contained a 175-nt hybrid exon obtained from joining the 5’ end of β globin exon 2 to the 3’ end of β globin exon 1 (Dominski and Kole 1991a; Modafferi and Black 1997). The minigene reporter contains an exonic and upstream intronic PTBP binding site from c-Src. The test exon is flanked by wild-type β globin exons 1 and 2 (Amir-Ahmady et al. 2005). The minigene Dup4-5 contains the Src N1 exon with upstream and downstream intronic regulatory regions inserted between the flanking β globin exons of Dup175 (Modafferi and Black 1999; Chou et al. 2000; Chan and Black 1995). The minigene Dup4-1 E8a contains the mouse E8a exon with two PTBP binding sites within the upstream intron inserted between the flanking β globin exons 1 and 2 (Tang et al. 2011). Multi-site RBD3, RBD4, and RBD3-4 glutamine and arginine mutants were created by GenScript.

### Reverse-transcription PCR

RNA was isolated from N2A cells using the PureLink RNA Mini Kit (Invitrogen) and reverse transcribed with oliog DT and Superscript III (Life Technologies) according to the manufacturer’s instructions. Spliced products were PCR amplified (16 cycles) using a 5’ primer (DUP-8) 5’-GACACCATGCATGGTGCACCTG-3’ and Cy3-labeled 3’ primer (DUP-3) 5’-AACAGCATCAGGAGTGGACAGATCCC-3’ (Integrated DNA Technologies). The PCR products were separated by 8% acrylamide/7.5 M urea denaturing gels and visualized by a Typhoon FLA7000 PhosphorImager. Bands corresponding to the length of included and excluded products were boxed and band intensity was quantified using ImageQuant TL software. Percent spliced-in values were calculated by dividing the band intensity of the included product by the total intensity of included and excluded products and multiplying by 100. Percent spliced-out values were calculated by dividing the band intensity of the excluded product by the total intensity of included and excluded products and multiplying by 100. Statistical significance was determined by one-way analysis of variance (ANOVA) with post hoc Tukey Honestly Significant Difference (HSD) test using Python. A p-value <0.05 was considered statistically significant. The boxplots were generated using R software (http://shiny.chemgrid.org/boxplotr/).

### Immunoblottin**g**

Whole-cell lysates were separated by 12% acrylamide SDS-PAGE, transferred to an Immobilon PVDF membrane, and probed with anti-Flag (Sigma catalog# F3165) and GAPDH antibodies (Invitrogen 6C5 via Fisher Scientific) at a 1:2500 dilution. After incubation with fluorescent-conjugated Alexa488 secondary antibody (Invitrogen A11001 via Fisher Scientific) at a 1:2500 dilution, the blots were scanned using a Typhoon FLA7000 PhosphorImager.

### Multiple Sequence Alignment and Web Logo

17 PTBP1 protein sequences were collected from different species, 15 from vertebrate and 2 from non-vertebrate species. The vertebrate species ranged from mammals to fish, and the non-vertebrate species included roundworm and fruit fly. These sequences were retrieved from NCBI (www.ncbi.nlm.nih.gov). The protein sequences were collected and aligned using Mega11 ClustalW. The amino acids that were fully conserved were highlighted in light blue and the lysine residues known to be acetylated were indicated by a black tick mark above. Dots represent identical amino acids. This multiple sequence alignment was then uploaded to UC Berkeley weblogo website (https://weblogo.berkeley.edu/logo.cgi) to create the weblogo. The weblogos were constrained to RRM3 and RRM4 of PTBP1. The size of each amino acid letter indicates the frequency the amino acid was observed among the 17 species chosen.

### Ion distribution calculations

PTBP RBD3 and RBD4 structures with and without bound RNA were prepared using structural data from the Protein Data Bank (PDB IDs: 2ADC and 2EVZ) (Oberstrass et al. 2005; Vitali et al. 2006). The first model from each PDB file was used. The 2EVZ structure was RMSD-aligned to 2ADC using cpptraj (Roe and Cheatham 2013)to ensure a consistent frame of reference. Histidine protonation states were retained as listed in the PDB files and were designated as HID in all cases. Force field parameters were assigned using the Amber ff19SB force field for proteins and OL3 for RNA (Pérez et al. 2007; Banáš et al. 2010; Kovalenko and Hirata 1999). Wild-type simulations used the structures and sequences directly from the 2ADC and 2EVZ files. To investigate the impact of lysine acetylation, we generated mutant structures, in which lysine residues at positions 394, 400, 424, 436, 454, 508, 511, and 515 were substituted with glutamine. These correspond to residues 368, 374, 398, 410, 428, 482, 485, and 489 in the 2ADC structure, and 45, 51, 75, 87, 105, 159, 162, and 166 in the 2EVZ structure.

Three-dimensional reference interaction site model (3D-RISM) calculations were performed using dielectrically consistent one-dimensional RISM (DRISM) calculations as input, with both methods implemented in AmberTools 24(Chandler and Andersen 1972; Hirata and Rossky 1981). The dielectric constant was set to 78.446, with a 0.2 M KCl concentration, and the bulk water concentration was 55.0 M. Ion parameters were taken from the Joung–Cheatham set and the cSPC/E water model was used (Joung and Cheatham 2008; Kovalenko et al. 2010). DRISM calculations were performed on a 32,768-point grid with 0.025 Å spacing, with a temperature of 298.15 K. For 3D-RISM, calculations used a buffer of 24 Å around the solute and a 0.25 Å grid spacing.

The partial series expansion of order-3 (PSE-3) closure relation was used, and convergence was defined as reaching a residual tolerance of 10⁻⁵ for 3D-RISM and 10⁻¹² for DRISM (Kast and Kloss 2008). All equations were solved using the MDIIS algorithm (Kovalenko, Ten-No, and Hirata 1999).

## Acknowledgements

This work was supported in part by NIH grant 1SC3GM132036 to N.M.K, a CSUBIOTECH Award to N.M.K and C.N. and the CSU Fullerton Office of Research and Sponsored Projects Jr/Sr award to N.M.K. T.L. was supported by the National Science Foundation (NSF) under Grants, MRI-2320718 and MRI-2320846.

